# *Leptospira interrogans* prevents macrophage cell death and pyroptotic IL1β release through its atypical lipopolysaccharide

**DOI:** 10.1101/2022.07.25.501344

**Authors:** Delphine Bonhomme, Veronica Hernandez-Trejo, Stylianos Papadopoulos, Rémi Pigache, Martine Fanton d’Andon, Ahmed Outlioua, Ivo G. Boneca, Catherine Werts

## Abstract

*Leptospira interrogans* are bacteria that can infect all vertebrates and are responsible for leptospirosis, a neglected zoonosis. Some hosts are susceptible to leptospirosis whereas mice are resistant and get chronically colonized. Although leptospires escape recognition by some immune receptors, they activate NLRP3-inflammasome and trigger IL1β secretion. Classically, IL1β secretion is associated with lytic inflammatory cell death called pyroptosis, resulting from cytosolic LPS binding to inflammatory caspases. Interestingly, we showed that *L. interrogans* do not trigger cell death in either murine, human, hamster, or bovine macrophages, escaping both pyroptosis and apoptosis. Strikingly, we also revealed in murine cells, a potent antagonistic effect of leptospires and their atypical LPS on spontaneous and *E. coli* LPS-induced cell death. The leptospiral LPS efficiently prevents caspase 11 dimerization and subsequent gasdermin D cleavage. Finally, we showed that pyroptosis escape by leptospires prevents massive IL1 β release, and we consistently found no major role of IL1-Receptor in controlling experimental leptospirosis *in vivo*. Overall, our findings described a novel mechanism by which leptospires dampen inflammation, thus potentially contributing to their stealthiness.

**Graphical abstract:** 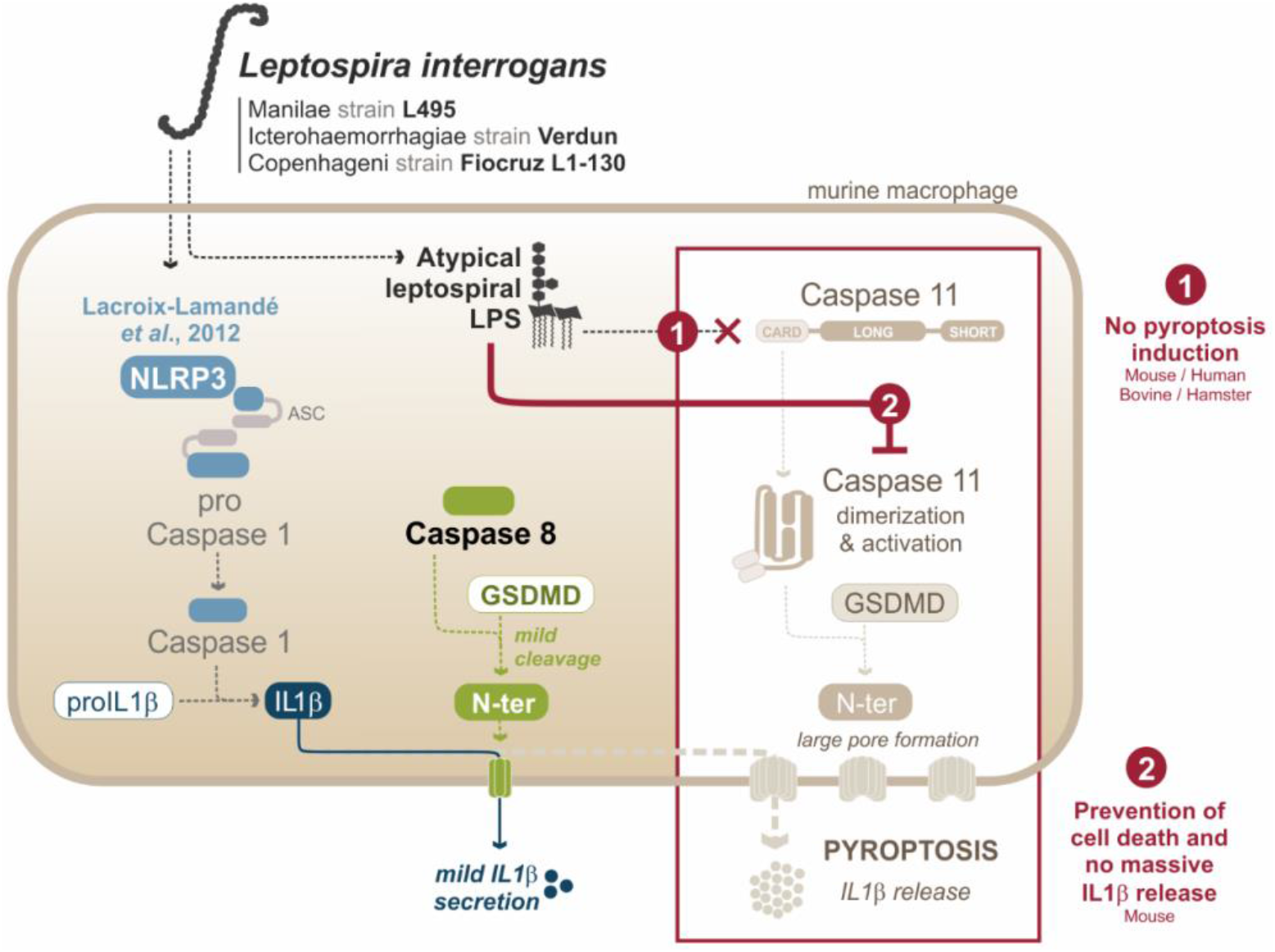

## Introduction

Leptospirosis is a re-emerging zoonosis caused by pathogenic spirochetes bacteria called leptospires. Among more than 60 species, *L. interrogans* is the most prevalent and pathogenic species. It can reside either free in the environment or infect vertebrates, although not all hosts show clinical signs of the disease. Humans are sensitive to leptospirosis and can present symptoms that vary from a mild flu-like syndrome to multi-organs failure, also known as Weil’s disease, lethal in 5 to 10 % cases (Adler, 2015; Costa *et al*, 2015). *L. interrogans* can also provoke acute symptoms in cattle, pigs and horses, such as morbidity, abortion, or uveitis (Adler, 2015). Among the resistant animals, mice and rats do not present symptoms of acute infection, however they get chronically colonized in the kidneys’ proximal tubules, leading to excretion of the bacteria in the urine, contributing to the zoonotic transmission of *L. interrogans* (Ko *et al*, 2009; Ratet *et al*, 2014).

Upon infection, innate immunity is the first line of defense that is activated to trigger inflammatory and antimicrobial responses. Recognition of pathogens occurs through Pattern-Recognition Receptors (PRRs), that have evolved to bind conserved molecules of microbes called Microbe-Associated Molecular Patterns (MAMPs), such as transmembrane Toll-Like Receptors (TLRs) and cytosolic NOD and NOD-Like Receptors (NLRs). Among them, NOD-Like Receptor Pyrin 3 (NLRP3) requires integration of two signals for activation. Priming of NLRP3 occurs by stimulation of other PRRs leading to NF-κB translocation, hence allowing NLRP3 and cytokines mRNA transcription (Bauernfeind *et al*, 2009; Franchi *et al*, 2009). NLRP3 is then activated by many cellular stresses, such as UV, crystals, Reactive Oxygen Species (ROS) potassium (K^+^) efflux or calcium (Ca^2+^) influx (Schroder & Tschopp, 2010). NLRP3 oligomerizes and binds ASC adaptors to form signaling platforms called canonical inflammasomes. Inflammasomes allow cleavage of caspase 1, a protease that, in turn, cleaves pro-Interleukin-1β (pro-IL1β) and pro-Interleukin-18 (pro-IL18) into mature cytokines, that are potent inflammatory mediators (Martinon *et al*, 2002; Wang *et al*, 2002). Furthermore, NLRP3 has also been associated with the activation of caspases 4-5 (human) / caspase 11 (mouse) both participating to the so-called “non-canonical inflammasome”, leading to inflammatory lytic cell death with release of the cytosolic content, called pyroptosis. Caspases 4-5/11 self-cleave upon intracellular binding of their CARD domain to the lipid A of intracellular LPS (Shi *et al*, 2014) and then trigger cleavage of gasdermin D (GSDMD) (Huang *et al*, 2019; Kayagaki *et al*, 2013; Shi *et al*., 2014). Subsequently, the N-terminal fragment of GSDMD oligomerizes to form large pores in the host cell membrane, leading to lytic cell death (Liu *et al*, 2016). NLRP3 hence contributes, through canonical and non-canonical inflammasomes, to the production and release of inflammatory cytokines in response to infection. However, the interplay between canonical and non-canonical inflammasomes remains unclear (Kayagaki *et al*., 2013).

Leptospires are stealth diderm bacteria. Their cell wall classically harbors peptidoglycan (PG), but is characterized by two endoflagella, and numerous tri-acylated lipoproteins. Furthermore, unlike *Borrelia burgdorferi* and *Treponema pallidum*, leptospires possess a lipopolysaccharide (LPS) in their outer membrane (Vinh *et al*, 1986). LPS is one of the few identified virulence factors (Murray *et al*, 2010) and allows classification of leptospires in more than 300 immunologically distinct serovars (Caimi & Ruybal, 2020; Ko *et al*., 2009). Several leptospiral mechanisms to escape recognition of MAMPs by macrophages have previously been described (Santecchia *et al*, 2020). The leptospiral PG escapes NOD1 and NOD2 recognition through tight association of PG with the LipL21 lipoprotein that prevents muropeptides release and NOD1/2 signaling (Ratet *et al*, 2017). The leptospiral endoflagella, because of its periplasmic localization, escapes TLR5 signaling (Holzapfel *et al*, 2020). Furthermore, the leptospiral LPS is structurally peculiar. Its lipidic anchor, also known as lipid A, possesses specific features: *(i)* unsaturated acyl chains, *(ii)* amine bonds and *(iii)* missing and methylated phosphate groups in position 4’ and 1, respectively (Que-Gewirth *et al*, 2004). Consequently, the leptospiral LPS escapes recognition by human TLR4, although it is recognized by murine TLR4 (Nahori *et al*, 2005). In addition, the carbohydrate component of the LPS, also known as O antigen, is more complex that the repeated sugar units present in *E. coli* or *S. enterica* LPS (Bonhomme & Werts, 2020; Cinco *et al*, 1986). Also, lipoproteins always co-purify with the leptospiral LPS, hence conferring its ability to signal through TLR2 (Werts *et al*, 2001). Both the O antigen and the co-purifying lipoproteins have been shown to play a role in the prevention of murine TLR4 endocytosis and escape from TLR4-TRIF endosomal signaling (Bonhomme *et al*, 2020). To our interest, leptospires have been shown to activate NLRP3 inflammasome in murine macrophages, with signal 1 being TLR2/TLR4 activation by the leptospiral LPS/lipoproteins and signal 2 being the downregulation of sodium/potassium pump by leptospiral glycolipoprotein (GLP) (Lacroix-Lamande *et al*, 2012). Such inflammasome activation triggers cleavage of caspase 1 and production of inflammatory IL1β (Lacroix-Lamande *et al*., 2012). Another study reported that leptospires also induce NLRP3 in human cells and induce IL1β and IL18 production (Li *et al*, 2021; Li *et al*, 2018). Interestingly, in human but not in murine macrophages, the activation of NLRP3 would be dependent on ROS production (Lacroix-Lamande *et al*., 2012; Li *et al*., 2021; Li *et al*., 2018). However, the activation of caspases 4-5/11 by the leptospiral LPS had never been investigated. Therefore, whether leptospires also trigger induction of non-canonical inflammasome and pyroptotic cell death or not remains unanswered.

Cell death of macrophages upon infection by leptospires has previously been investigated (Santecchia *et al*., 2020). Although no study specifically assessed pyroptosis induction, some studies reported induction of apoptosis through caspases 8 and 3 in monkey and mouse cell lines (Jin *et al*, 2009; Merien *et al*, 1997). However, using different leptospiral strains, other studies report no cytotoxicity (Toma *et al*, 2011). Interestingly, we recently showed that leptospires actively enter and exit macrophages, without replicating, and with no loss of cell viability (Santecchia I *et al*, 2022). It was therefore to our interest to address the potential toxicity of different leptospiral pathogenic strains in primary macrophages derived from mouse bone marrow. Furthermore, because of the ability of *L. interrogans* to infect other hosts such as humans, bovines and hamsters; and because several species-specific mechanisms were discovered in the innate immune escape of leptospires (Holzapfel *et al*., 2020; Nahori *et al*., 2005), we also addressed the potential cytotoxicity of leptospires on human THP1-CD14 monocytes and on bone marrow derived macrophages from calf and hamsters

Overall, our study aimed at characterizing potential cytotoxic effects upon infection by *L. interrogans* and to assess whether leptospires induce the non-canonical inflammasome in addition to the canonical NLRP3 in cells from both resistant and sensitive hosts. Our results suggest that leptospires do not trigger the non-canonical inflammasome. We further focused on the role played by the peculiar leptospiral LPS in the lack of induction and inhibition of cell death.

## Results

### Although they induce IL1β secretion in murine BMDMs, leptospires prevent cell death

Our group previously demonstrated that infection by *L. interrogans* serovar Copenhageni strain Fiocruz L1-130 triggers NLRP3 inflammasome, induces caspase 1 cleavage and triggers IL1β production in murine BMDMs (Lacroix-Lamande *et al*., 2012). We first assessed whether such mechanism was conserved for other strains of *L. interrogans*. BMDMs were infected for 24 h with three frequently studied pathogenic serovars of *L. interrogans* (serovar Manilae strain L495, serovar Icterohaemorrhagiae strain Verdun and serovar Copenhageni strain Fiocruz L1-130). First, Western blot analyses showed that the three strains of leptospires similarly induce the cleavage of caspase 1 in its active form of p10 subunit (**Figure 1A**). Consistently, all strains induced in a dose dependent manner the secretion of IL1β, measured in the supernatants of RAW-ASC cells (a murine macrophage cell line with a functional inflammasome) (**Figure 1B**). Considering the activation of the canonical inflammasome by the three pathogenic serovars of *L. interrogans*, we then investigated whether leptospires also trigger the activation of the non-canonical inflammasome and hence induce pyroptosis. As a marker of cell membrane damage and lytic cell death, we measured the leakage of lactate dehydrogenase (LDH), a constitutive cytosolic enzyme. LPS from *E. coli* with ATP (inflammasome inducer) was used as a positive control of pyroptotic cell death. Unexpectedly, although leptospires activate the canonical inflammasome, they did not induce LDH release in the supernatant of BMDMs 24h post-infection (**Figure 1C**), even at high MOI, suggesting that no membrane damage is occurring. Surprisingly, compared with the level of spontaneous LDH released by non-stimulated cells, we even observed a reduction of the LDH released upon infection with the three strains of leptospires and in the same range of magnitude at all MOIs. Interestingly, such phenotype was recapitulated upon stimulation with either heat-killed leptospires or purified LPS (**Figure 1D**). Using MTT assay as a measure of cellular metabolic activity, we confirmed that the viability of the macrophages was not reduced upon infection with leptospires (**Figure 1E**), and we even observed higher MTT levels upon infection. Similar results were obtained upon stimulation by heat-killed bacteria or LPS (**Figure 1F**). Altogether, these results show that leptospires are not cytotoxic for BMDMs and suggest that they can even prevent spontaneous cell death. Of note, LDH and MTT reflect membrane integrity and metabolism, respectively, and their dosage does not exactly reflect the number of BMDMs. Therefore, to conclude on a potential effect on cell death prevention, we enumerated by high-content (HC) microscopy the number of BMDMs at the time of plating and 24h later, with or without leptospiral infection or stimulation with their LPS. A small reduction in the number of adherent cells was observed in the non-infected condition (**Figure 1G**), most likely due to basal spontaneous death. Surprisingly, infection with leptospires or stimulation with their LPS efficiently prevented this spontaneous decrease, which was not observed with *E. coli* LPS (**Figure 1G**), here showing that leptospires not only are not cytotoxic, but also prevent BMDMs cell death.

**Figure 1.**
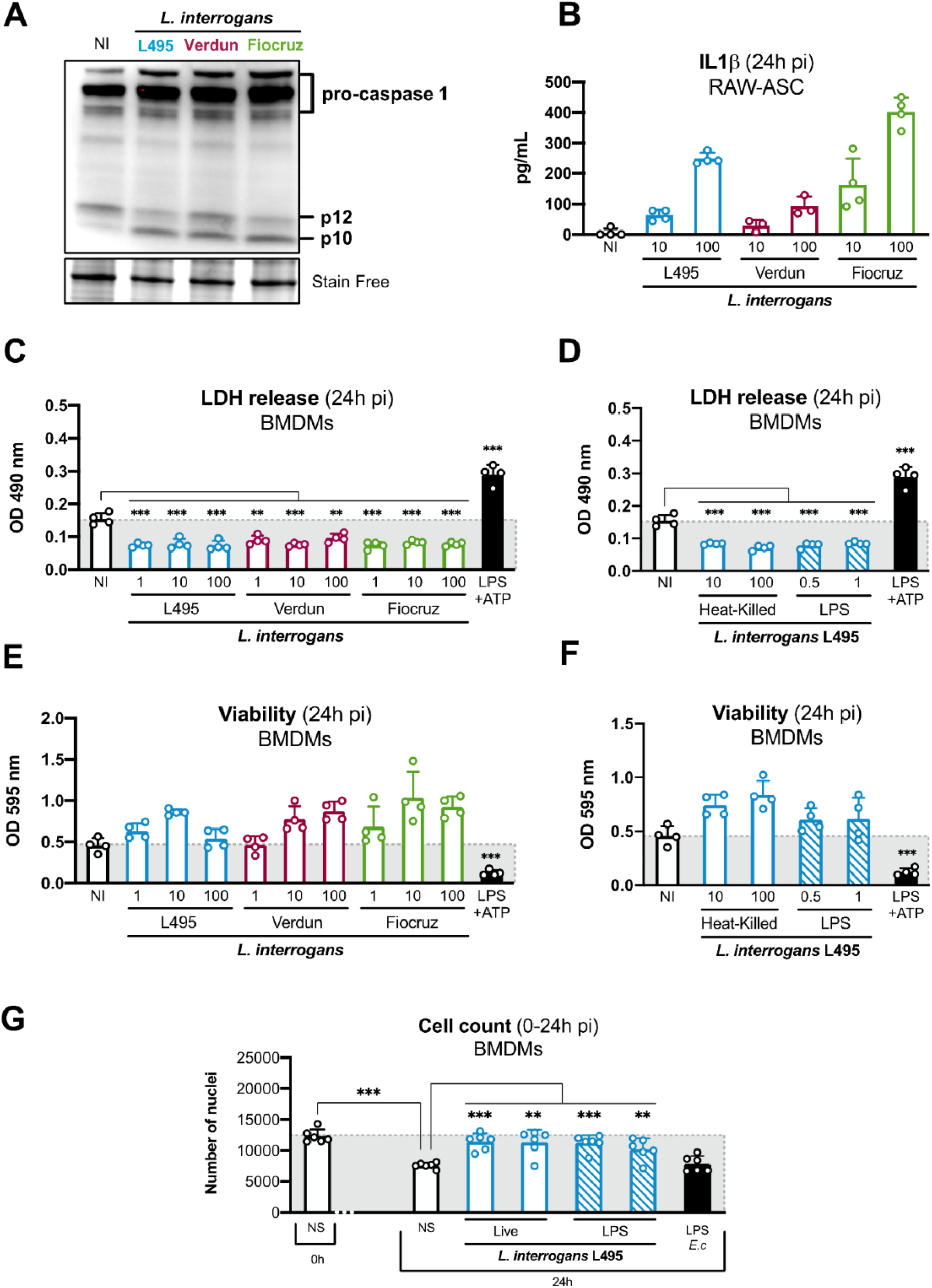
Although they induce IL1β secretion in murine BMDMs, leptospires prevent cell death. **A)** Western Blot analysis of caspase 1 in BMDMs after 8h infection with MOI 100 of *L. interrogans* serovar Manilae strain L495 (blue), serovar Icterohaemorrhagiae strain Verdun (red), serovar Copenhageni strain Fiocruz L1-130 (green). **B)** IL1β dosage by ELISA in the supernatant of RAW-ASC cells after 24h infection with MOI 10-100 of the three serotypes of *L. interrogans* mentioned above. **C-D)** LDH release measured by CyQuant assay on the supernatant of BMDMs after 24h infection with **C)** MOI 1-100 of the three serotypes of *L. interrogans* mentioned above. **D)** MOI 10-100 of heat-killed strain L495 or 24h stimulation with 0.5-1 μg/mL of purified leptospiral LPS from strain L495. **E-F)** Cell viability measured by MTT assay in BMDMs after 24h infection with **E)** MOI 1-100 of the three serotypes of *L. interrogans* mentioned above, **F)** MOI 10-100 of heat-killed strain L495 or 24h stimulation with 0.5-1 μg/mL of purified leptospiral LPS from L495. **C-E)** Positive control is 1 μg/mL of *E. coli* LPS + 5 mM ATP. **G)** BMDMs enumeration by high-content (HC) microscopy at the time of cell plating (0h), or after 24h with either infection by MOI 10-100 of *L. interrogans* serovar Manilae strain L495 or stimulation with 0.1-1 μg/mL of its purified LPS. Control stimulation is performed with 1 μg/mL of *E. coli* LPS. **B-G)** Bars correspond to mean +/− SD of technical replicates (*n*>4). **A-G)** Data presented are representative of at least 3 independent experiments.

### Leptospires do not trigger the molecular pathways of apoptosis or pyroptosis in murine BMDMs

Since infection with *L. interrogans* did not induce LDH release nor alter macrophage viability, we specifically investigated the apoptotic and pyroptotic molecular pathways by studying the cleavage of caspase 3 and caspase 11 / GSDMD, respectively. Consistent with the absence of cytotoxicity, leptospires did not trigger caspase 3 cleavage, after overnight infection (**Figure 2A**), and no caspase 3/7 activity was measured 24h post-infection, in contrast to staurosporine, the positive control of apoptosis. These results showed that leptospires do not trigger apoptosis in murine BMDMs. Interestingly, caspase 3/7 activity basal level was even reduced in BMDMs infection with leptospires, here again suggesting that leptospires might prevent cell death. Regarding pyroptosis, Western blot analyses of caspase 11 at 8h post-infection revealed that infection with *L. interrogans* serovar Manilae strain L495 induced upregulation of caspase 11, both at the mRNA and protein levels, like *E. coli* LPS + ATP, the positive control for inflammasome activation that induced complete cell death with 5 mM ATP, and only partial cell death at 2 mM (**Figure 2B**). Since the caspase 11 antibody did not recognize the p25 cleaved form of the caspase, we could not conclude regarding its potential cleavage. However, the parallel study of GSDMD by Western blot showed that infection with leptospires only triggers minor GSDMD cleavage and does not induce massive accumulation of GSDMD-N-ter, the moiety forming pores, as observed in pyroptotic cells, and confirmed by quantification (**Figure 2C**). Overall, these findings show that leptospires do not trigger the apoptotic and pyroptotic molecular pathways, confirming the lack of cytotoxicity observed beforehand.

**Figure 2.**
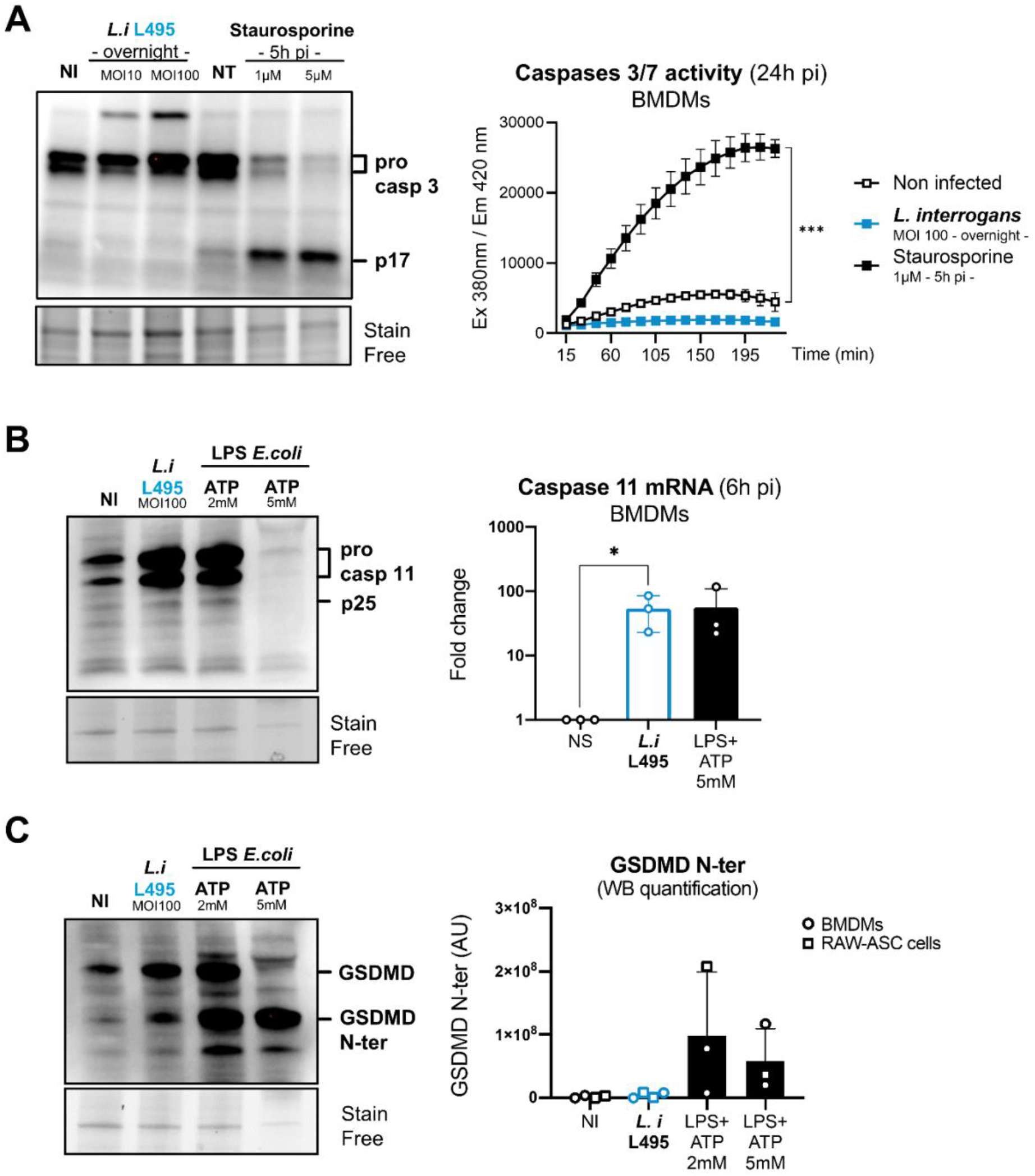
Leptospires do not trigger apoptosis or pyroptosis molecular pathways in murine BMDMs. **A)** *Left panel*. Western Blot analysis of caspase 3 in BMDMs after overnight infection with MOI 10-100 of *L. interrogans* serovar Manilae strain L495 or control stimulation during 5h with 1-5 μM of staurosporine. **A)** *Right panel*. Kinetics of caspase 3/7 activity, assessed by fluorometry measures every 15 min of cleaved substrate Ac-D EVD-AMC, in BMDMs after overnight infection with MOI 100 of *L. interrogans* serovar Manilae strain L495, or control stimulation during 5h with 1 μM of staurosporine. Dots correspond to mean +/− SD of technical replicates (*n*=6). **B)** Western Blot and mRNA RT-qPCR analyses of caspase 11 in BMDMs after either 8h (WB, *left panel*) or 6h (qPCR, *right panel*) infection with MOI 100 of *L. interrogans* serovar Manilae strain L495 or control stimulation 1 μg/mL of *E. coli* LPS + 2-5 mM ATP. **C)** Western Blot analysis (*left panel*) and quantification of GSDMD (*right panel*) in BMDMs and RAW-ASC cells after 8h infection with MOI 100 of *L. interrogans* serovar Manilae strain L495 or control stimulation 1 μg/mL of *E. coli* LPS + 2-5 mM ATP. **B-C)** Bars correspond to the mean of at least three independent experiments. **A-C)** Data presented are representative of at least 3 independent experiments.

### Caspase 8 contributes to mild GSDMD cleavage and IL1β secretion upon leptospiral infection

Upon infection with leptospires, GSDMD is mildly cleaved, but does not lead to the lytic pyroptotic cell death. To the extent of our knowledge, GSDMD can be cleaved either by caspase 11 or, to a lower extent by caspase 1. Recent studies have also demonstrated that caspase 8 can cleave GSDMD (Chen *et al*, 2019; Orning *et al*, 2018; Sarhan *et al*, 2018). In addition to caspase 1 activation (Lacroix-Lamande *et al*., 2012), leptospires were also shown to activate caspase 8 in macrophages (Jin *et al*., 2009). Therefore, to address the role of these different caspases in the mild GSDMD cleavage observed upon infection, we infected RAW-ASC cells in presence of different caspase inhibitors and observed 24h post-infection the GSDMD cleavage by Western blot. The use of the caspase 8 inhibitor and of the pan-caspases inhibitor, but not the caspase 1/11 inhibitor, led to a reduction of the faint GSDMD-N-ter band induced upon *Leptospira* infection (**Figure 3A and Figure 3B**), showing that caspase 8 contributes to the mild GSDMD cleavage upon infection. After cleavage, the N-terminal fragment of GSDMD classically accumulates in the cell to form small pores, and then large pores, that can lead to mild IL1 β secretion and lytic cell death, respectively (Heilig *et al*, 2018). To address whether GSDMD cleavage by caspase 8 could induce small pores playing a role in IL1β secretion, we measured the cytokine in the supernatant of cells treated with the different caspase inhibitors. Inhibiting caspase 1/11 resulted in lower IL1β secretion, as expected considering that caspase 1 is cleaving the pro-IL1β into the mature cytokine (**Figure 3C**). Interestingly, we also observed a reduction in IL1β secretion when using the caspase 8 inhibitor (**Figure 3C**), suggesting that GSDMD cleavage by caspase 8 contributes to the mild IL1β secretion observed upon infection. The efficiency of the caspase 8 and pan-caspases inhibitors was controlled using staurosporine, an apoptosis inducer, and by monitoring in Western blot the reduction in the p18 cleaved form of caspase 8 (**Figure 3D**).

**Figure 3.**
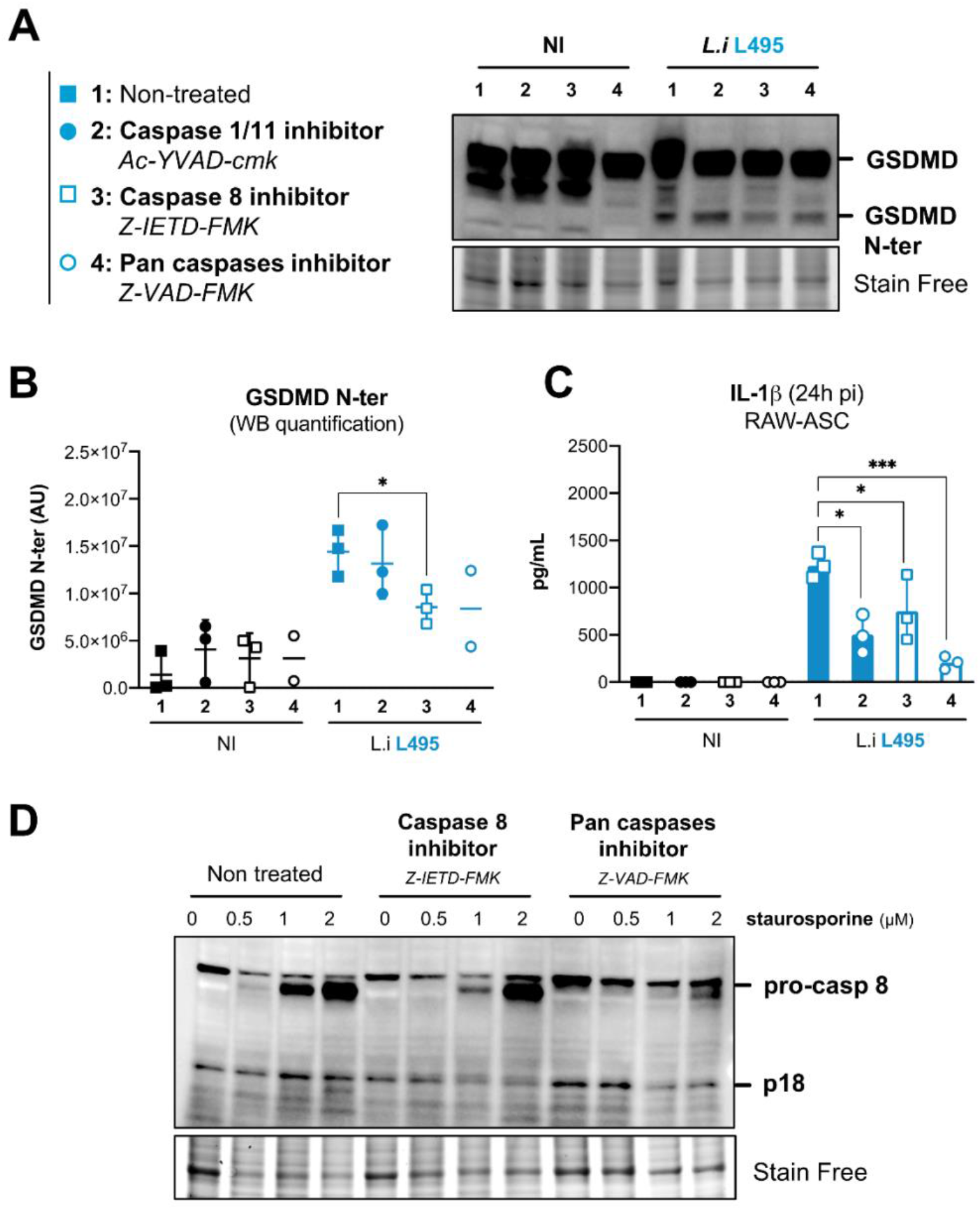
Caspase 8 contributes to mild GSDMD cleavage and IL1β secretion upon leptospiral infection. Western Blot analysis and **B**) quantification of GSDMD in RAW-ASC and **C)** IL1β dosage by ELISA in the supernatant of cells pretreated for 30 min with the caspase 1/11 inhibitor (Ac-YVAD-cmk, 50 μM), caspase 8 inhibitor (Z-IETD-FMK, 20 μM) or pan-caspase inhibitor (Z-VAD-FMK, 10 μg/mL) and infected for 16h with MOI 100 of *L. interrogans* serovar Manilae strain L495. Bars correspond to mean +/− SD of technical replicates (*n*=3). **D)** Western Blot analysis of caspase 8 in RAW-ASC cells pretreated with caspase 8 inhibitor (Z-IETD-FMK, 20 μM) or pan-caspase inhibitor (Z-VAD-FMK, 10 μg/mL) and stimulated for 5h with 0.5-2 μM of staurosporine. **A-D)** Data presented are representative of at least 3 independent experiments.

### Transfected leptospiral LPS escapes pyroptosis and even prevents spontaneous cell death

Considering that caspase 1 and caspase 11 did not contribute to the mild GSDMD cleavage observed upon infection and considering that no cell death was observed, we hypothesized that *L. interrogans* does not trigger the non-canonical inflammasome. Classically, it is the binding of cytosolic *E. coli* LPS (lipid A) to caspase 11 (CARD domain) that leads to GSDMD cleavage and pyroptosis [36]. To test whether the LPS of *L. interrogans* would also fulfill this function, purified leptospiral LPS was transfected into RAW-ASC cells and we monitored pyroptotic cell death. First, we addressed 6h post transfection, by epifluorescence microscopy, the morphological properties of the transfected cells, identifiable *via* GFP fluorescence, produced from a plasmid co-transfected with the LPS. As expected, cell transfected (Empty) without LPS displayed physiological morphology, whereas cell transfected with *E. coli* LPS were loosely adherent with damaged membranes as observed in bright field, and no longer labeled with GFP (**Figure 4A**). Of note, reduction of the GFP signal in these cells suggests that membrane permeability was altered, allowing for the GFP signal leakage, consistent with the induction of pyroptosis. In contrast, cells transfected with the leptospiral LPS showed similar features as the negative control cells, suggesting that no pyroptosis had occurred (**Figure 4A**). To further confirm this results, transfected cells were stained with propidium iodide (PI), a fluorescent dye staining nuclei that only enters in cells upon rupture of the plasma membrane. Cells were then analyzed by flow cytometry (**Figure 4B**) and fluorimetry (**Figure 4C**) 12h and 24h post-transfection, respectively. Both methods confirmed that the LPS of *L. interrogans* did not induce PI accumulation even at 24h post-transfection, unlike the classical LPS of *E. coli*, that efficiently triggered pyroptosis. Interestingly, the levels of PI fluorescence were even lower upon transfection of the leptospiral LPS than in the negative control cells, suggesting that the leptospiral LPS contributes to cell death prevention. Furthermore, LDH release was monitored 24h post-transfection with different amounts of leptospiral LPS from *L. interrogans* serovar Manilae strain L495, serovar Icterohaemorrhagiae strain Verdun and serovar Copenhageni strain Fiocruz L1 130. We evidenced no LDH release upon transfection with LPS of any of the three pathogenic serovars (**Figure 4D**). Interestingly here again we observed that leptospires, through their LPS, reduced the basal level of spontaneous LDH release of murine macrophages. Finally, to confirm that the leptospiral LPS did not trigger the non-canonical inflammasome, we performed native gel analysis of caspase 11 (known to dimerize upon activation) after LPS transfection. Upon transfection of *E. coli* LPS, a shift of the band that could correspond to the dimerized caspase 11 was visible as expected (**Figure 4E**). However, this shift was not observed upon transfection of the leptospiral LPS (**Figure 4E**). Overall, these data show that transfected leptospiral LPS does not induce caspase 11 dimerization and subsequent pyroptotic cell death thus escaping activation of the non-canonical inflammasome.

**Figure 4.**
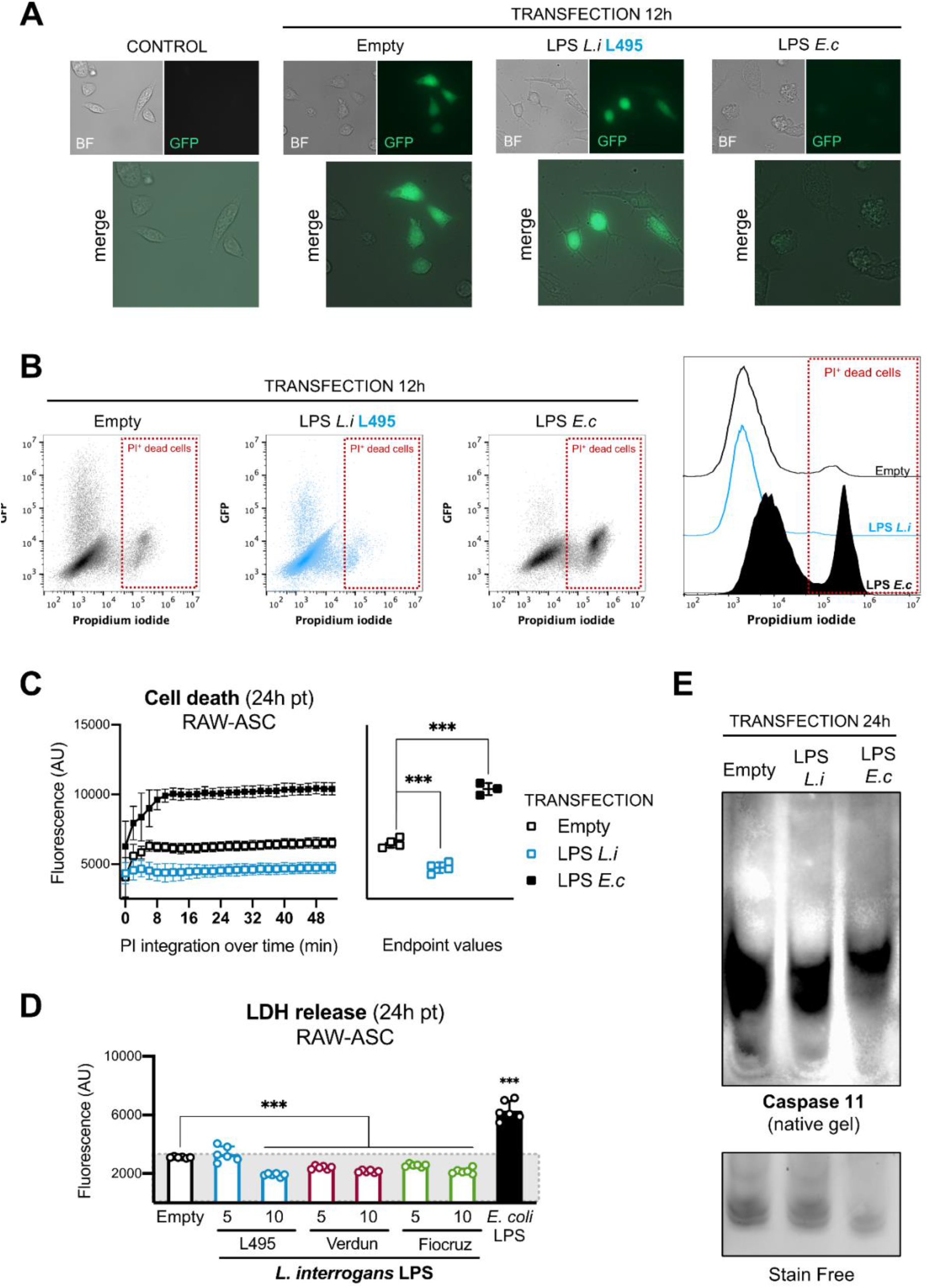
Transfected leptospiral LPS escapes pyroptosis and even prevents spontaneous cell death. **A)** Epifluorescence microscopy of RAW-ASC cells after 6h transfection with pCMV-GFP and either 10 μg/mL *E. coli* LPS or 10 μg/mL *L. interrogans* LPS from serovar Manilae strain L495. **B)** Cell death monitored by flow cytometry analysis of propidium iodide (PI) integration in RAW-ASC cells after 12h transfection with pCMV-GFP and either 10 μg/mL *E. coli* LPS or 10 μg/mL *L. interrogans* LPS from serovar Manilae strain L495**. C)** Cell death monitored by fluorimetry analysis of PI integration in RAW-ASC cells after 24h transfection with pcDNA3 and either 10 μg/mL *E. coli* LPS or 10 μg/mL *L. interrogans* LPS from serovar Manilae strain L495. Lines correspond to mean +/− SD of technical replicates (*n*=4). **D)** LDH release measured by CyQuant assay on the supernatants of RAW-ASC cells after 24h transfection with pcDNA3 and either 10 μg/mL *E. coli* LPS or 10 μg/mL *L. interrogans* LPS from serovar Manilae strain L495 (blue), serovar Icterohaemorrhagiae strain Verdun (red) or serovar Copenhageni strain Fiocruz L1-130 (green). Bars correspond to mean +/− SD of technical replicates (*n*=6). **E)** Western Blot analysis in native conditions of caspase 11 in RAW-ASC cells after 24h transfection with pCMV-GFP and either 10 μg/mL *E. coli* LPS or 10 μg/mL *L. interrogans* LPS from serovar Manilae strain L495**. A-E)** Data presented are representative of at least 3 independent experiments.

### Leptospiral LPS potently inhibits *E. coli* LPS-induced cell death

Our results showed that the leptospiral LPS could prevent macrophage cell death by reducing spontaneous LDH release and PI accumulation upon transfection. We therefore investigated if the leptospiral LPS could also protect against a strong induction of pyroptosis induced by *E. coli* LPS. To address this, we compared cell transfected with either *L. interrogans* LPS, *E. coli* LPS or cells co-transfected with equal amounts of both LPS. Strikingly, the leptospiral LPS was able to strongly reduce both PI accumulation (measured by cytometry and fluorimetry) and LDH release upon *E. coli* LPS exposure (**Figure 5A**). Interestingly, we also observed that LPS from *L. interrogans* serovar Manilae strain L495, serovar Icterohaemorrhagiae strain Verdun and serovar Copenhageni strain Fiocruz L1-130 all decreased LDH release upon *E. coli* LPS exposure, although LPS from L495 was more potent (**Figure 5A,** *right panel*). Furthermore, we addressed at a molecular level both the dimerization of caspase 11 and the cleavage of GSDMD upon LPS transfection. Interestingly, the LPS of *L. interrogans* led to a reduction of both caspase 11 dimerization and GSDMD cleavage upon exposure to LPS of *E. coli* (**Figure 5B**). Classically, the mechanism by which *E. coli* LPS activates caspase 11 involves direct binding of the lipid A to the CARD domain of the enzyme, hence triggering its dimerization. Therefore, we decided to analyze potential binding of *L. interrogans* LPS and *E. coli* LPS to recombinant caspase 11. However, we only found commercial recombinant caspase 11 enzyme that lacked the CARD binding domain (**Figure 5C**). Nevertheless, we incubated this recombinant caspase 11 with the different LPS and analyzed their migration on highly reticulated polyacrylamide gels (20%) that do not allow the migration of the LPS / bound enzyme, but only allow migration of the unbound material. As expected, the *E. coli* LPS did not bind the enzyme lacking its CARD domain. Interestingly, we observed a binding of the leptospiral LPS to this recombinant caspase 11 (**Figure 5D**). Although we could not address the specificity nor the nature of the interaction, this result suggests an atypical interaction between the leptospiral LPS and caspase 11, that could play a role in the cell death inhibition. Finally, as we observed that the leptospiral LPS was strongly inhibiting cell death by targeting caspase 11 of the non-canonical inflammasome, we excluded a role for the canonical inflammasome by showing that this inhibitory effect was conserved in both RAW-ASC and RAW264.7 cells that respectively harbor and lack a functional canonical inflammasome (**Figure 5E**).

**Figure 5.**
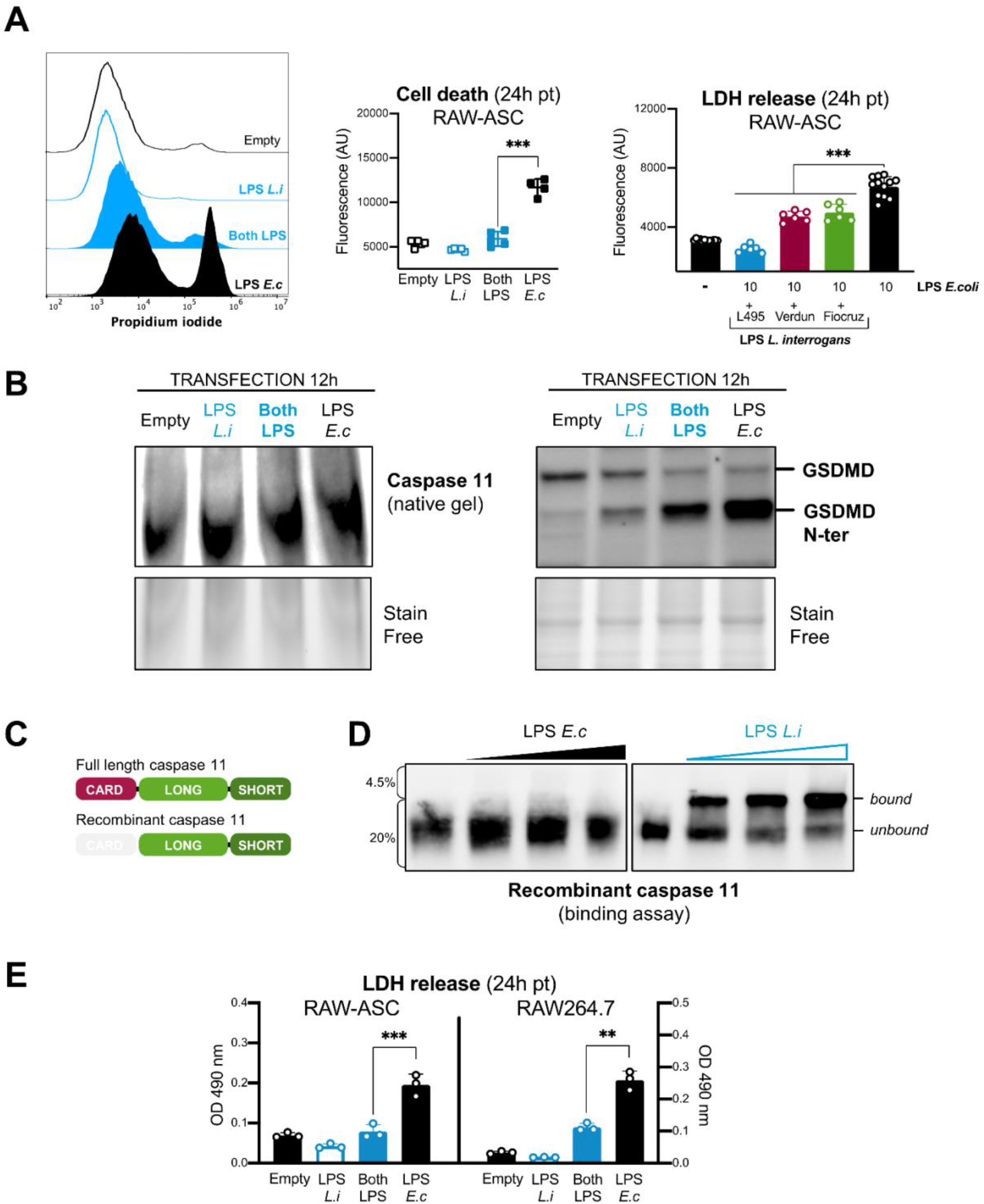
Leptospiral LPS potently inhibits *E. coli* LPS-induced cell death. **A)** *Left panels*. Cell death monitored by flow cytometry and fluorimetry analysis of PI integration in RAW-ASC cells, 12h and 24h, respectively, after transfection with pCMV-GFP and 10 μg/mL *E. coli* LPS, 10 μg/mL *L. interrogans* LPS from serovar Manilae strain L495 or 10 μg/mL of both LPS. **A)** *Right panel*. LDH release measured by CyQuant assay on the supernatant of RAW-ASC cells after 24h transfection with pcDNA3 and 10 μg/mL *E. coli* LPS + 10 μg/mL *L. interrogans* LPS from serovar Manilae strain L495 (blue), serovar Icterohaemorrhagiae strain Verdun (red) or serovar Copenhageni strain Fiocruz L1-130 (green). Bars correspond to mean +/− SD of technical replicates (*n*=6). **B)** *Left panel*. Western Blot analysis in native conditions of caspase 11 in RAW-ASC cells after 12h transfection with pCMV-GFP and 10 μg/mL *E. coli* LPS, 10 μg/mL *L. interrogans* LPS from serovar Manilae strain L495 or 10 μg/mL of both LPS. **B)** *Right panel*. Western Blot analysis of GSDMD in RAW-ASC cells after 12h transfection with pCMV-GFP and 10 μg/mL *E. coli* LPS, 10 μg/mL *L. interrogans* LPS from serovar Manilae strain L495 or 10 μg/mL of both LPS. **C)** Schematic representation of full length and recombinant caspase 11 lacking the LPS-binding CARD domain. **D)** Binding assay between recombinant caspase 11 and *E. coli* LPS or *L. interrogans* LPS from serovar Manilae strain L495, performed on highly reticulated 20% polyacrylamide native gels that do not allow the entrance of LPS and bound material and only allow migration of unbound proteins. **E)** LDH release measured by CyQuant assay on the supernatant of either RAW-ASC or parental RAW264.7 cells 24 h after transfection with pcDNA3 and 10 μg/mL *E. coli* LPS, 10 μg/mL *L. interrogans* LPS from serovar Manilae strain L495 or 10 μg/mL of both LPS. Bars correspond to mean +/− SD of technical replicates (*n*=3). **A-E)** Data presented are representative of at least 3 independent experiments.

### Leptospires do not trigger pyroptosis in human THP1-CD14 cells

After establishing that leptospires are not cytotoxic for mouse macrophages, we further investigated potential cytotoxicity on cells from other hosts, namely humans that are susceptible to leptospirosis. To do so, we infected human monocyte-like THP1 cells stably transfected with CD14 (hereafter called THP1-CD14) with the main three pathogenic serovars of *L. interrogans* (serovar Manilae strain L495, serovar Icterohaemorrhagiae strain Verdun and serovar Copenhageni strain Fiocruz L1-130). After 24 h of infection, we controlled that the strains induced the production of IL1β in THP1-CD14, illustrating the activation of the canonical inflammasome (**Figure 6A**). We then addressed the key features of the non-canonical inflammasome. GSDMD analysis by Western blot revealed no visible cleavage of the protein (**Figure 6B**) and we observed no LDH release nor viability decrease upon infection with any of the leptospiral strains (**Figure 6C**). Consistent with the results obtained in murine BMDMs, these data overall show that, although they induce IL1 β secretion, leptospires do not trigger pyroptosis in human THP1-CD14 cells either.

**Figure 6.**
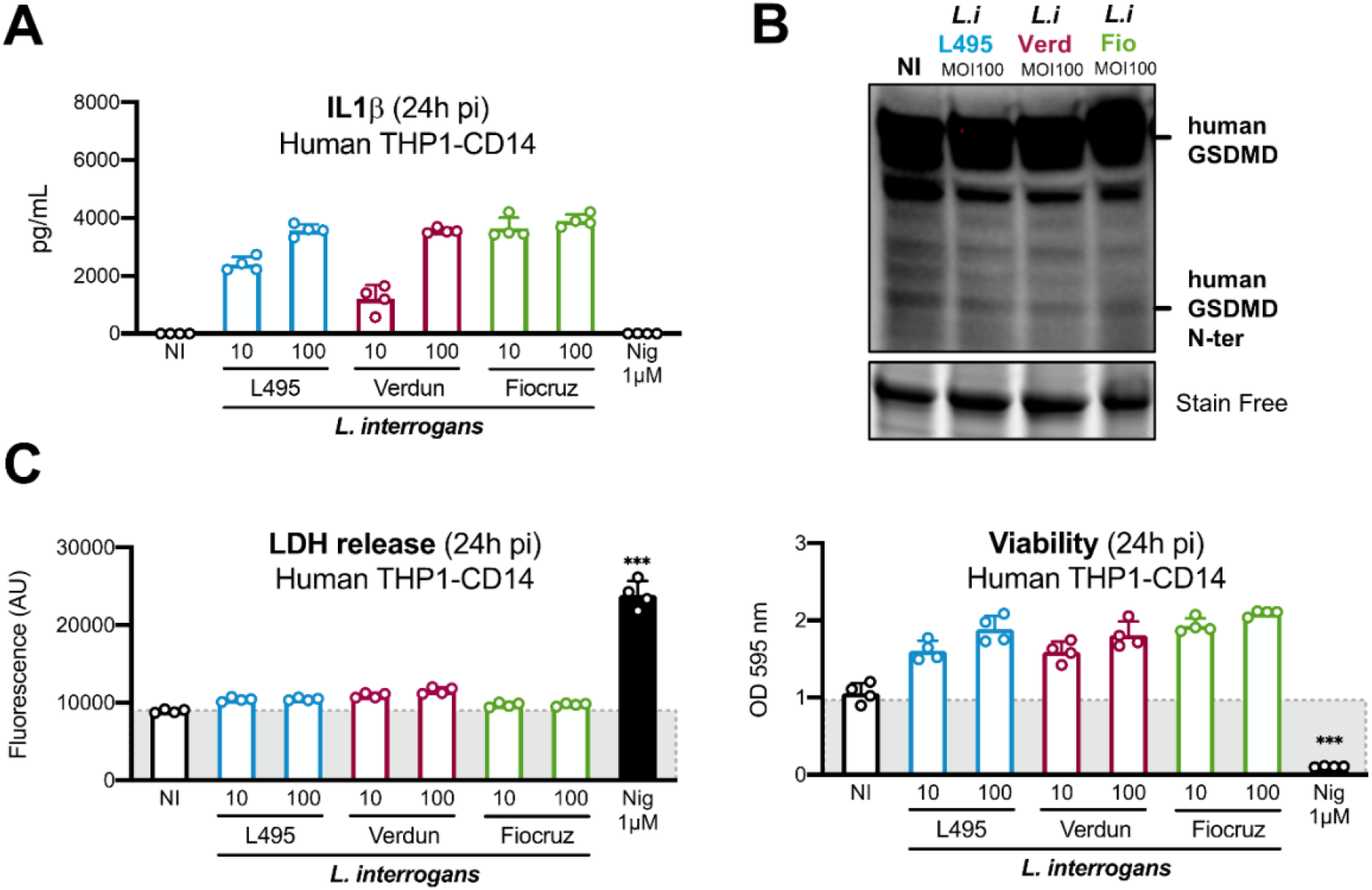
Leptospires do not trigger pyroptosis in human THP1-CD14 cells. **A)** IL1β dosage by ELISA in the supernatant of human THP1-CD14 cells 24h after infection with MOI 10-100 of *L. interrogans* serovar Manilae strain L495 (blue), serovar Icterohaemorrhagiae strain Verdun (red) or serovar Copenhageni strain Fiocruz L1-130 (green). **B)** Western Blot analysis of human GSDMD in THP1-CD14 after 16h infection with MOI 100 of the three serotypes of *L. interrogans* mentioned above. **C)** LDH release measured by CyQuant assay and cell viability measured by MTT assay on human THP1-CD14 cells after 24h infection with MOI 10-100 of the three serotypes of *L. interrogans* mentioned above. Positive control is 1 μM of nigericin. Bars correspond to mean +/− SD of technical replicates (*n*=4). **A-C)** Data presented are representative of at least 3 independent experiments.

### Leptospires do not trigger pyroptosis in bovine and hamster BMDMs

To study the potential induction of pyroptosis on primary cells from susceptible hosts, we derived BMDMs from bone marrow of calf and hamster. These cells were then infected with the main three pathogenic serovars of *L. interrogans* for 24 h (serovar Manilae strain L495, serovar Icterohaemorrhagiae strain Verdun and serovar Copenhageni strain Fiocruz L1-130). First, we observed no visible cell damage on the infected cells, unlike cells treated with inflammasome induced LPS + ATP (**Figure 7A and Figure 7B, upper panels**). Viability and LDH assays further showed no cell death and no membrane alteration, respectively (**Figure 7A and Figure 7B, middle panels**). Finally, as a control of infection, we measured the production of nitric oxide. Results show that both bovine BMDMs and hamster BMDMs are efficiently stimulated by leptospires and produce NO in response to the infection (**Figure 7A and Figure 7B, lower panels**).

**Figure 7.**
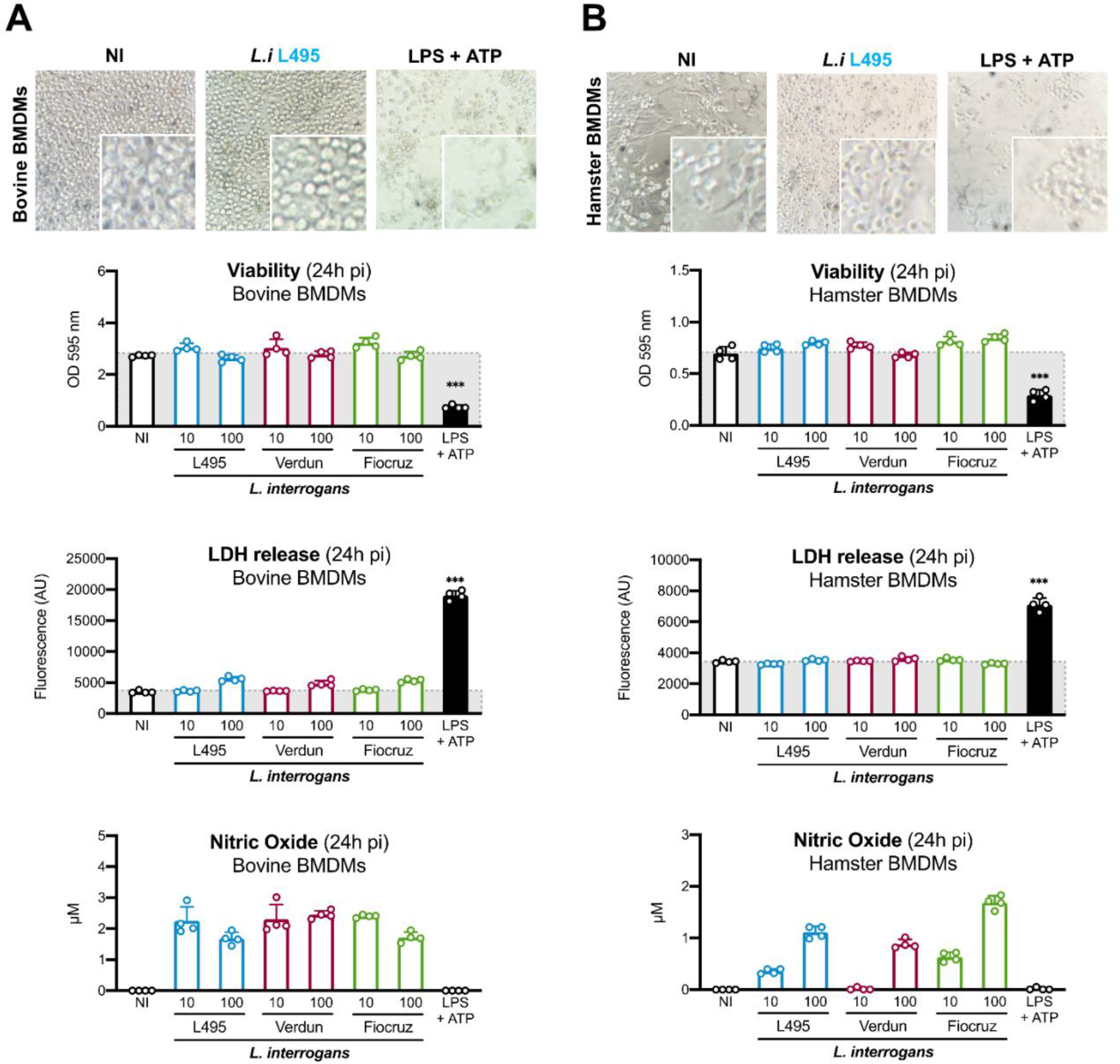
Leptospires do not trigger pyroptosis in bovine and hamster BMDMs. **A-B)** Microscopy images, cell viability (measured by MTT assay), LDH release (measured by CyQuant assay), and nitric oxide production (measured by Griess reaction) on **A)** bovine and **B)** hamster BMDMs 24h after infection with MOI 10-100 of *L. interrogans* serovar Manilae strain L495 (blue), serovar Icterohaemorrhagiae strain Verdun (red) or serovar Copenhageni strain Fiocruz L1-130 (green). Positive control of pyroptosis is 1 μg/mL of *E. coli* LPS + 5 mM ATP. Bars correspond to mean +/− SD of technical replicates (*n*=4). Data presented are representative of **A)** 3 independent experiments or **B)** 2 independent experiments.

### Pyroptosis escape by leptospires prevents massive IL1β release and IL1R signaling pathway does not contribute to the control of experimental leptospirosis

Lastly, we investigated the functional consequences of the escape of pyroptosis by leptospires. We hypothesized that, leptospiral escape of pyroptosis greatly dampens the IL1β secretion induced upon infection. To test such hypothesis *in vitro*, we infected RAW-ASC cells with *L. interrogans* and artificially induced pyroptosis by adding ATP 3 h before measuring IL1 β in cell supernatants. Results showed that leptospires alone induced only mild IL1β production 24h post infection. Interestingly, upon infection followed by ATP treatment, the levels of IL1β in the supernatant were 5 times higher (**Figure 8A, left panel**). This massive release was further shown to be specific to IL1β considering that the levels of the chemokine RANTES were not modified with or without the ATP trigger of cell pyroptosis (**Figure 8A, right panel**). Furthermore, considering that leptospires do escape massive IL1β secretion *in vitro*, we decided to further test the importance of the IL1 β signaling pathway *in vivo* on experimental leptospirosis. Leptospirosis is a biphasic disease that comprises an initial acute blood replication of leptospires that, in resistant mice, disappear from the blood stream before reappearing around 8 days post-infection localized in the proximal tubules of the kidneys, where they establish a life-long chronic colonization (Ratet *et al*., 2014). Depending on the bacterial inoculum at the time of infection, experimental leptospirosis using mouse models allows the study of both the acute and the chronic stages. To address the role of ILR in experimental leptospirosis, we therefore performed 2 infections of WT and IL1Receptor (IL1R) knock-out (KO) mice with *L. interrogans* serovar Copenhageni strain Fiocruz L1-130. The first infection was performed with a high leptospiral inoculum (2×10^8^ bacteria/mouse), to study the role of ILR on a severe acute model (Ratet *et al*., 2014). Inflammation was monitored in the kidneys at the acute phase of the disease, 3 days post-infection. Interestingly, no statistically significant differences were observed in either RANTES, IL6 or IL1 β between WT and IL1R KO mice infected with leptospires (**Figure 8B, upper panel**). The second infection was performed with a lower leptospiral inoculum (2×10^6^ bacteria/mouse), to study the role of ILR in the chronic model of leptospirosis. The leptospiral loads associated with the chronic kidney colonization were measured by qPCR and showed no statistically significant differences in the loads of the WT and ILR KO mice (**Figure 8B, lower panel**). Overall, these results show that ILR signaling does not majorly contribute to the control of leptospirosis, either at the acute or chronic phase of the disease.

**Figure 8.**
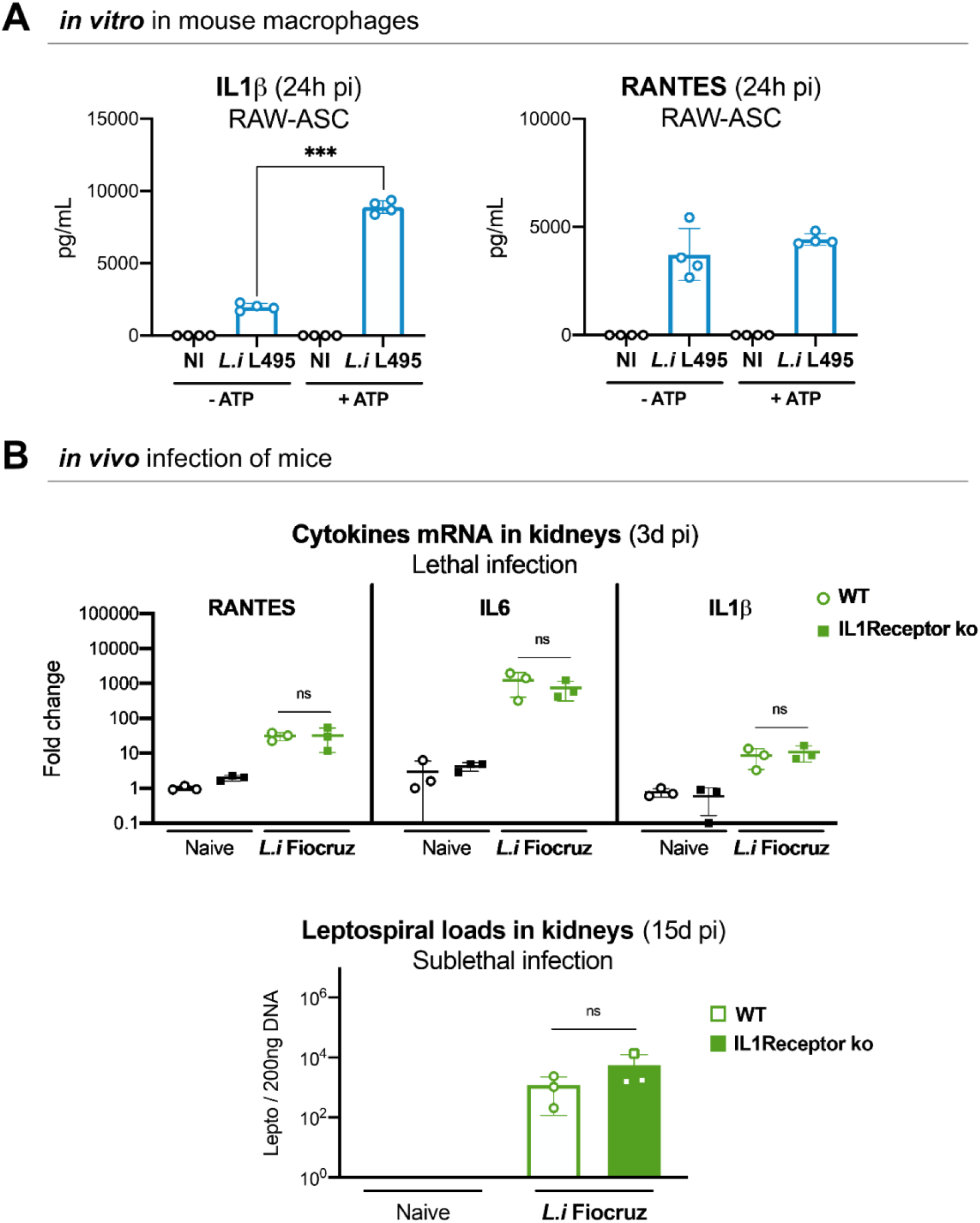
Pyroptosis escape by leptospires prevents massive IL1β release and IL1R signaling pathway does not contribute to the control of experimental leptospirosis. **A)** IL1β and RANTES dosage by ELISA in the supernatant of RAW-ASC cells 24h after infection with MOI 100 of *L. interrogans* serovar Manilae strain L495, with or without addition of 5 mM ATP 3h before supernatant collection. Bars correspond to mean +/− SD of technical replicates (*n*=4). Data presented are representative of 3 independent experiments. **B)** *Upper panel*. RANTES, IL6 and IL1β mRNA levels measured by RT-qPCR in kidneys 3 days post intraperitoneal infection with 2×10^8^ *L. interrogans* serovar Copenhageni strain Fiocruz L1-130 of either WT C57BL/6J or IL1Receptor (IL1R) knock-out (ko) mice. Bars correspond to mean +/− SD of individual animals (*n*=3). **B)** *Lower panel*. Leptospiral loads measured by qPCR in kidneys 15 days post intraperitoneal infection with 2×10^6^ *L. interrogans* serovar Copenhageni strain Fiocruz L1-130 of either WT C57BL/6J or IL1Receptor (IL1R) knock-out (ko) mice. Bars correspond to mean +/− SD of individual animals (*n*=3).

## Discussion

Our results showed that infection with the three of the main pathogenic serovars of *L. interrogans* (serovars Manilae, Copenhageni and Icterohaemorrhagiae) did not induce cell death in murine, bovine and hamster BMDMs, nor in human THP1-CD14 monocyte-like cells. Considering that leptospirosis is a zoonosis that affects differently all vertebrates, and because the species-specificity of the innate immune response might play an essential role in the course of the infection, it was our interest to address leptospiral cytotoxicity in different hosts. Mice are resistant to the acute form of the disease and get chronically infected in their kidneys. On the other hand, humans and bovines are susceptible and present diverse symptoms (Bonhomme & Werts, 2022), and hamsters are very susceptible to the disease and are the referent model for acute human leptospirosis (Gomes *et al*, 2018). We therefore showed that the lack of induced macrophage cell death upon infection with leptospires is a conserved cellular mechanism in both susceptible and resistant hosts. In murine BMDMs, we further excluded induction of both pyroptosis and apoptosis, with no cleavage of GSDMD and caspase 3, respectively. However, in the current literature, it has been described that *L. interrogans* serovar Icterohaemorrhagiae strain Verdun induces apoptosis in Vero cells and mouse J774 macrophages (Merien *et al*., 1997) and that other serovars of *L. interrogans* could kill macrophages through either necrosis (necroptosis/pyroptosis) (Hu *et al*, 2013) or through induction of caspase 8/3-dependent apoptosis (Jin *et al*., 2009). In contrast, another publication using *L. interrogans* serovar Manilae (Toma *et al*., 2011) confirms that infection does not have major cytotoxic effects on murine macrophages. To reconcile the current literature with our results, we therefore hypothesize that different serovars of *L. interrogans* and/or infection of different cell lines could differentially induce cell death. We are also aware that cell culture conditions, especially over confluency of the cells at the time of infection with leptospires, induce biased and aspecific cell death that could partially explain the discrepant results in the literature.

To the extent of our knowledge, *L. interrogans* would be the first bacteria to efficiently trigger the canonical NLRP3 inflammasome (Lacroix-Lamande *et al*., 2012), and yet escape subsequent induction of pyroptosis. Furthermore, although the IL1β production triggered by leptospires is mediated by NLRP3 and caspase 1, its release upon infection seems to be atypical. Indeed, our results showed that leptospires only triggered mild GSDMD cleavage, independent of both caspase 1 and caspase 11, but dependent of caspase 8. Several recent studies have highlighted the versatility of caspase 8, usually involved in apoptosis, but also involved in numerous cell death pathways (Fritsch *et al*, 2019; Han *et al*, 2021; Orning *et al*., 2018). Indeed, few studies reported GSDMD cleavage and pyroptosis induction by caspase 8 (Chen *et al*., 2019; Orning *et al*., 2018; Sarhan *et al*., 2018). However, we do not know how leptospiral-induced GSDMD cleavage is maintained at non-lytic levels. Therefore, further studies are required to understand the underlying mechanism of this non-lethal role of caspase 8. We hypothesize, as suggested (Heilig *et al*., 2018), that limited accumulation of GSDMD N-ter fragments could lead to the formation of small non-lytic pores, that would only allow mild secretion of IL1β.

Classically, human caspases 4-5 and murine caspases 11 dimerize and are activated upon binding of the lipid A moiety of the cytosolic LPS through their CARD domain (Ross *et al*, 2018; Shi *et al*., 2014). Our results showed that transfection of the leptospiral LPS did not induce the dimerization of caspase 11, unlike LPS of *E. coli*, suggesting that the activation of the enzyme did not occur. It remains to be determined what leptospiral LPS feature is responsible for this phenotype. Our results showed that different leptospiral serovars, defined by their O antigen (Cinco *et al*., 1986; Patra *et al*, 2015; Vinh *et al*., 1986) and that harbor a conserved lipid A (Eshghi *et al*, 2015; Novak *et al*, 2022; Que-Gewirth *et al*., 2004), all escape pyroptosis induction to a similar extent, suggesting that the leptospiral lipid A is responsible for the phenotype. Caspases 4-5/11 all respond to both penta- and hexa-acylated lipids A but tetra-acylated lipids A are not recognized by murine caspase 11 (Hagar *et al*, 2013; Lagrange *et al*, 2018). For instance, LPS of *Francisella novicida* is tetra-acylated, and several other tetra-acylated LPS (*Helicobacter pylori, Rhizobium galegae*) have been shown to escape caspase 11 induction (Kayagaki *et al*., 2013), suggesting that penta or hexa-acylation levels are prerequisite for caspase 11 activation (Zamyatina & Heine, 2020, 2021). Considering that the leptospiral lipid A is hexa-acylated, our main hypothesis is therefore that the other features of the lipid A (methylated 1-position phosphate group and lacking 4’-position phosphate group, unsaturated acyl chains and amide liaisons) (Que-Gewirth *et al*., 2004) could be responsible for lack of interaction with caspase 11 and pyroptosis escape. Such hypothesis would be consistent with a study suggesting that lack 4’-position phosphate group in *F. novicida* lipid A would also contribute to caspase 11 induction escape (Hagar *et al*., 2013; Zamyatina & Heine, 2020). Unfortunately, we were not able to directly assess the binding of the leptospiral lipid A to the CARD domain of caspase 11, given that commercial recombinant inflammatory caspases are devoid of their CARD domains, most probably because of their activation and cleavage upon binding of endogenous endotoxins present during the expression process. A study consistently reported that production of caspase 11 in *E. coli* is not appropriate for the study of non-activated caspase 11(Shi *et al*., 2014).

In addition to the pyroptosis escape mechanism of the leptospiral LPS, our results strikingly evidenced an antagonistic effect of the leptospiral LPS on spontaneous cell death and furthermore on *E. coli* LPS induced cell death. Inhibitory effects on pyroptosis induction have previously been described for the atypical LPS of *Helicobacter pylori* and *Rhizobium galegae*. However, in these cases, the mechanism is linked to the tetra-acylation of the lipid A (Kayagaki *et al*., 2013). For *L. interrogans*, our results interestingly showed that the LPS of different serovars, that have conserved lipid A (Eshghi *et al*., 2015; Novak *et al*., 2022; Que-Gewirth *et al*., 2004), but that differ in the structure of their O antigen (Cinco *et al*., 1986; Patra *et al*., 2015; Vinh *et al*., 1986) do not have the same ability to inhibit *E. coli* LPS induced cell death. We therefore favor a role for the O antigen in the inhibitory effect of the leptospiral LPS. Interestingly, the carbohydrate component of the leptospiral LPS is peculiar and does not have the same sugar composition than the repeated units of classical LPS such as the one of *E. coli* (Bonhomme & Werts, 2020; Cinco *et al*., 1986; Patra *et al*., 2015; Vinh *et al*., 1986). Although the O antigen section of the LPS is not supposed to be involved in caspase 11 activation, several studies previously reported that it could interfere with activation of the non-canonical inflammasome. *Salmonella* and *Shigella* both trigger low pyroptosis whereas mutant strains with shorter or completely lacking O antigen induce higher levels of pyroptosis (Mylona *et al*, 2021; Watson *et al*, 2019). It was believed that the full-length O antigen could interfere with the lipid A binding to caspase 11 (Mylona *et al*., 2021). Our results showing that the leptospiral LPS binds caspase 11 through an atypical interaction, independent of the CARD domain, further support the hypothesis that steric hindrance could prevents caspase 11 activation by *E. coli* LPS and consequent pyroptosis induction. Overall, we believe that the escape and antagonistic effects of the leptospiral LPS would be linked to two different mechanisms, involving the lipid A and the O antigen, respectively.

On the host side, we showed that the antagonistic effect of leptospiral LPS was independent of the ASC adaptor and consequently independent of the canonical-inflammasome, suggesting involvement of components of the non-canonical inflammasome. Our results further demonstrated that the leptospiral LPS was able to prevent caspase 11 dimerization, showing that the inhibitory effect occurred upstream of the caspase 11-GSDMD-pyroptosis pathway. We therefore propose that the leptospiral LPS, through its atypical features, prevents pyroptosis very early on, which could explain the striking efficiency of the inhibition. Of note, this could also partially explain why a multiplicity of infection of only 1 bacterium *per* macrophage is able to efficiently inhibit spontaneous cell death. Interestingly, other pyroptosis prevention mechanisms occur downstream of caspase 11, with direct GSDMD targeting (Kang *et al*, 2018). It is the case of disulfiram, an FDA-approved drug, that covalently modifies cysteines 191/192 in GSDMD and prevents pore formation (Hu *et al*, 2020).

This study is consistent with previous studies from our laboratory showing that leptospires already escape NOD1/2, TLR5 and TLR4-TRIF responses in murine macrophages, hence considerably dampening the production of cytokines, chemokines, and antimicrobial compounds (Bonhomme *et al*., 2020; Holzapfel *et al*., 2020; Ratet *et al*., 2017) making leptospires very discrete bacteria. We had previously showed that leptospires activate NLRP3 inflammasome and trigger the production of IL1β (Λαχρoιξ–Λαμαvδε *ετ αλ*., 2012). Classically, the IL1 β secreted by macrophages allows, once systemic, to induce fever and hepatic responses, such as C-reactive protein and complement response. Furthermore, IL1 β induces potent amplification of inflammation through IL1 receptor (IL1R) (Medzhitov, 2010). Our results showed that pyroptosis escape upon leptospiral infection results in concealing IL1β inside macrophages. In addition, our *in vivo* results showed that the course of leptospirosis is similar in WT and IL1R knock-out (KO) mice and suggest that the mild IL1 β secretion induced upon infection is not enough to trigger a strong inflammatory response.

Overall, our study revealed a novel immune escape mechanism by which leptospires decorrelate the canonical and non-canonical inflammasomes thus reducing IL1 β-mediated inflammation despite NLPR3 activation.

## Materials and Methods

### *Leptospira* strains and cultures

The pathogenic *L. interrogans* serovar Manilae strain L495, serovar Icterohaemorrhagiae strain Verdun and serovar Copenhageni strain Fiocruz L1-130 were grown in liquid Ellinghausen-McCullough-Johnson-Harris medium (EMJH) at 30 °C, without agitation. They were weekly passaged to be maintained in exponential phase. Bacteria were harvested for infection at the end of the exponential phase. Bacterial concentration was adjusted by centrifuging the culture at 4000 g for 25 minutes at room temperature. Bacteria resuspended in PBS were counted using a Petroff-Hauser chamber. Leptospires were also inactivated by heating at 56 °C for 30 min, under 300 rpm agitation.

### Purification of leptospiral LPS

LPS from the different leptospiral strains was purified using the hot water/phenol extraction method, as we recently reviewed (Bonhomme & Werts, 2020). Purified LPS was used for *in vitro* cell stimulation between 100 ng/mL and 10 μg/mL.

### Murine macrophages culture and infection

Bone marrow derived macrophages (BMDMs) were obtained after euthanasia by cervical dislocation of adult C57BL/6J mice. Isolation of the femurs, tibia and iliac bones was performed. Bone marrow cells were recovered by flushing out the bone marrow using a 22 G needle in complete RPMI medium (RMPIc): RPMI (Lonza) with 10% v/v heat inactivated foetal calf serum (HI-FCS, Gibco), 1 mM sodium pyruvate and 1X non-essential amino acids (Gibco) supplemented with 1X penicillin-streptomycin (Gibco). Erythrocytes were lysed using red blood cells lysis buffer (Sigma) and subsequent centrifugation at 1200 g for 7 minutes. Fresh or frozen bone marrow cells were differentiated into macrophages by seeding 5 x 10^6^ cells in cell culture dish (TPP) with 12 mL of RMPIc, supplemented with 1 X penicillin-streptomycin and 10 % v/v of L929 cells supernatant. Differentiation was carried out during 7 days with addition of 3 mL of medium at day 3. BMDMs were recovered at day 7 by scraping in PBS-EDTA 10 mM (Gibco) and centrifuged before counting and plating. Macrophage-like cell lines RAW-264.7 and RAW-ASC (Invivogen), cultivated in antibiotic-free RPMIc (10% v/v HI-FCS), were also used in this study. For all cells, plating was performed 24 hours before infection in antibiotic-free RPMIc (10% v/v HI-FCS) medium at a concentration of 0.3 x 10^6^ cells/mL for cell lines and 0.8 x 106 cells/mL for BMDMs. Cells were then infected in RPMIc (10% v/v HI-FCS) with live or heat-inactivated *Leptospira* strains or stimulated for 24 hours with leptospiral LPS (0.1-1 μg/mL), or with 1μg/mL *E. coli* LPS (Invivogen) and 2-5 mM ATP (Sigma). When indicated, RAW-ASC cells were pretreated 1h at 37°C before infection with the following caspase inhibitors: 50 μM Ac-YVAD-cmk, 20 μM Z-IETD-FMK or 10 mg/mL Z-VAD-FMK (Invivogen).

### Human THP1-CD14 cells culture and infection

Human THP1 monocyte-like cell line, stably transfected with CD14 (originally provided by Dr. Richard Ulevitch, Scripps, San Diego, CA, USA) were cultivated in antibiotic-free RPMIc medium (10% v/v HI-FCS). Cells were plated in RPMIc medium (2% v/v HI-FCS) at a concentration of 0.8 x 106 cells/mL 24h before infection. Cells were then infected in antibiotic-free RPMIc (2% v/v HI-FCS) with live *Leptospira* strains or stimulated for 24 hours nigericin (0.1-1μM).

### Bovine & hamster macrophages culture and infection

Bovine bone marrow cells were obtained from the femur of a neonate calf (kindly provided by Dr. Sonia Lacroix-Lamandé, INRA, Nouzilly, France). Hamster bone marrow cells were obtained from femurs and tibias of golden Syrian hamsters (kindly provided by Dr. Nadia Benaroudj, Institut Pasteur, Paris, France). In both cases, bone marrow cells were recovered by flushing out the bone marrow using RPMIc medium (20% v/v HI-FCS). Erythrocytes were lysed using red blood cells lysis buffer (Sigma) and subsequent centrifugation at 1200 g for 7 minutes. Frozen bone marrow cells were thawed and differentiated into macrophages by seeding 10 x 10^6^ cells in cell culture dish (TPP) with 12 mL of RPMIc (20% v/v HI-FCS), 1X penicillin-streptomycin (Gibco) and 5 ng/mL of recombinant human macrophage colony stimulating factor (hCSF, Peprotech). Differentiation was carried out during 14 days with addition of 3 mL of medium every 3 days. BMDMs were recovered at day 14 by scraping in PBS-EDTA 10 mM (Gibco) and centrifuged before counting. Cells were plated in RPMIc medium (20% v/v HI-FCS) at a concentration of 0.8 x 10^6^ cells/mL 24h before infection. Cells were then infected in antibiotic-free RPMIc (10% v/v HI-FCS) with live *Leptospira* strains or stimulated for 24 hours with 1μg/mL *E. coli* LPS (Invivogen) and 2-5 mM ATP (Sigma).

### MTT viability assay

MTT assay was performed on murine, bovine, hamster BMDMs, and on human THP1-CD14 cells 24h post-infection. Briefly, all the culture supernatant was removed and a solution of 1 mM MTT (Sigma) in RPMIc was added on the cells. After 90 min incubation at 37°C, dissolution of the formazan crystals was performed with isopropanol acidified with 1 M HCl. Optical density was measured at 595 nm on a BioTek ELx800 microplate reader.

### Cytosolic LDH release assay

LDH release was quantified on fresh cell culture supernatant of murine, bovine, hamster BMDMs, and on RAW-ASC, RAW264.7, THP1-CD14 cells 24h post-infection or 24h post-transfection of LPS. CyQuant LDH Cytotoxicity Colorimetric & Fluorimetric assays (Invitrogen) were performed according to the manufacturer’s instructions. Optical density was measured at 490 nm on a BioTek ELx800 microplate reader.

### Caspase 3/7 activity assay

Caspase activity was measured on murine BMDMs after overnight infection or 5 h stimulation with 1 μM of staurosporine (CST). Caspase 3/7 activity assay (CST) was performed according to the manufacturer’s instructions. Ac-DEVD-AMC substrate cleavage was immediately monitored by fluorescence at 380 nm (*ex*) / 420 nm (*em*) on a TECAN Spark fluorimeter (Life Sciences).

### Cytokine dosage by enzyme-linked immunosorbent assays (ELISA)

The secretion of cytokines (murine RANTES, murine IL1β and human IL1β) was assessed using cell culture supernatants of murine BMDMs, RAW-ASC and THP1-CD14 cells 24h post-infection. Supernatants were kept at −20 °C before cytokine dosage. ELISA assays (R&D DuoSet) were performed according to the manufacturer’s instructions. Optical density was measured at 450 nm on a BioTek ELx800 microplate reader, and cytokine concentration was determined using standard range provided by the manufacturer.

### Nitric oxide (NO) dosage by Griess reaction

NO dosage was performed on fresh cell culture supernatant of bovine and hamster BMDMs 24h post-infection. Briefly, Griess reaction was performed by incubation of 50 μL of fresh supernatant with 50 μL of 80 mM sulfanilamide in 2 M chlorohydric acid. Addition of 50 μL of 4 mM N-1-napthylethylenediamine dihydrochloride then allowed colorimetric determination of NO concentration. Optical density was measured at 540 nm on a BioTek ELx800 microplate reader and NO concentration was determined using standard range of nitrites.

### LPS transfection in RAW-ASC cells

For LPS transfection, RAW-ASC cells were plated in antibiotic-free RPMIc medium (10% v/v HI-FCS) at a concentration of 0.2 x 10^6^ cells/mL 24h before transfection, in transparent bottom black plates (Greiner) when fluorescence measurements where performed. Medium was removed and cells were transferred in FCS-free essential medium OptiMEM (Gibco) 2 h before transfection. Then, cells were transfected using FUGENE (Promega) in OptiMEM (Gibco), with the equivalent of 0.5 μL/well of FUGENE, 100 ng/well of DNA (either pcDNA3 or pCMV-GFP for fluorescence analyses) and LPS at a final concentration of 10 μg/mL. Quantities indicated correspond to the transfection of 96-well plates and were modified accordingly for transfection of 24-well plates. After either 6, 12 or 24 h of transfection, cells were analyzed by epifluorescence microscopy, flow cytometry or fluorimetry, respectively.

### Propidium iodide integration analysis by flow cytometry and fluorimetry

After either 12 or 24 h of transfection with LPS, RAW-ASC cells were stained with 1 mg/mL of propidium iodide (PI, Sigma). For flow cytometry analyses at 12 h, cells were then washed once in PBS with 2 mM EDTA and 0.5% v/v HI-FCS, scrapped and analyzed directly by flow cytometry on CytoFLEX (Beckman Coulter). Between 15 000 and 30 000 events were acquired for each condition and the data was analyzed with FlowJo V10 software. For fluorimetry analyses at 24 h, the kinetics of PI integration was monitored immediately at 535 nm (*ex*) / 617 nm (*em*) on a TECAN Spark fluorimeter (Life Sciences).

### Microscopy analyses

Live RAW-ASC cells transfected 6 h with LPS and fluorescent pCMV-GFP plasmid were observed under epifluorescent inverted microscope AxioObserver Z1 (Zeiss) with 63X oil-immersion objective. Brightfield illumination was maintained < 3V, laser intensity was maintained <50% and exposure time was automatically set from the brightest sample (empty transfection). Images analyses were performed using Fiji software. On the other hand, murine BMDMs enumeration was performed after 10 min fixation with 4% w/v *para*formaldehyde in PBS and 10 min staining with 1 μg/mL DAPI in PBS. After three washed with PBS, fixed cells were imaged on Opera Phenix HCS (Perkin Elmer) in confocal mode with 63X water-immersion objective. Laser intensity was maintained <25% and exposure time was set manually to obtain arbitrary unit (AU) of fluorescence around 1 000. Acquisition was performed automatically and >500 cells were analyzed *per* well. Image analysis was performed automatically using Colombus software (Perkin Elmer) and the number of nuclei in each condition was exported as output.

### SDS-PAGE and Western blots (WB)

Murine BMDMs, RAW-ASC and THP1-CD14 cells were collected 8-24 h post-infection or post-transfection by scrapping and centrifuged for 10 min at 1200 g. Cells were then lysed in 50 μL of RIPA lysis buffer (50 mM Tris, pH 7.5, 150 mM NaCl, 1% v/v Triton X-100, 0.5% w/v deoxycholate, 0.1% w/v SDS) supplemented with 1X complete Mini, EDTA-free protease inhibition cocktail (Roche) for 15 min on ice. After centrifugation for 30 min at full speed at 4°C, proteins were recovered in the supernatant and protein concentration was determined and adjusted by Bradford dosage (BioRad). Laemmli sample buffer, supplemented with 10% v/v βmercapto-ethanol, was added and samples were denatured at 99 °C for 10 min. SDS-PAGE of samples was performed on polyacrylamide 4-15% gradient gel in 1X Tris-Glycine-SDS buffer, at 110 V for 1 h. Internal loading control was performed by stain free visualization after 5 min activation on a ChemiDoc imaging system (BioRad). Protein transfer was done either on nitrocellulose membrane or polyvinylidene fluoride (PVDF) membrane (**Table 1**). Membranes were then saturated in TBS with 0.01% v/v Tween (TBST) with 5% w/v BSA for 1 h at room temperature. Incubation with primary antibodies was performed in TBST with 1% w/v BSA overnight at 4°C (**Table 1**). After three washes in TBST, secondary antibody incubation was done in TBST with 5% w/v BSA for 1h at room temperature under mild agitation (**Table 1**). After three washes in TBST, HRP activity was revealed using the SuperSignal West Femto Commercial TMB Substrate (ThermoFisher) on ChemiDoc imaging system (BioRad). WB quantification was performed on raw data in ImageLab software.

**Table 1.**
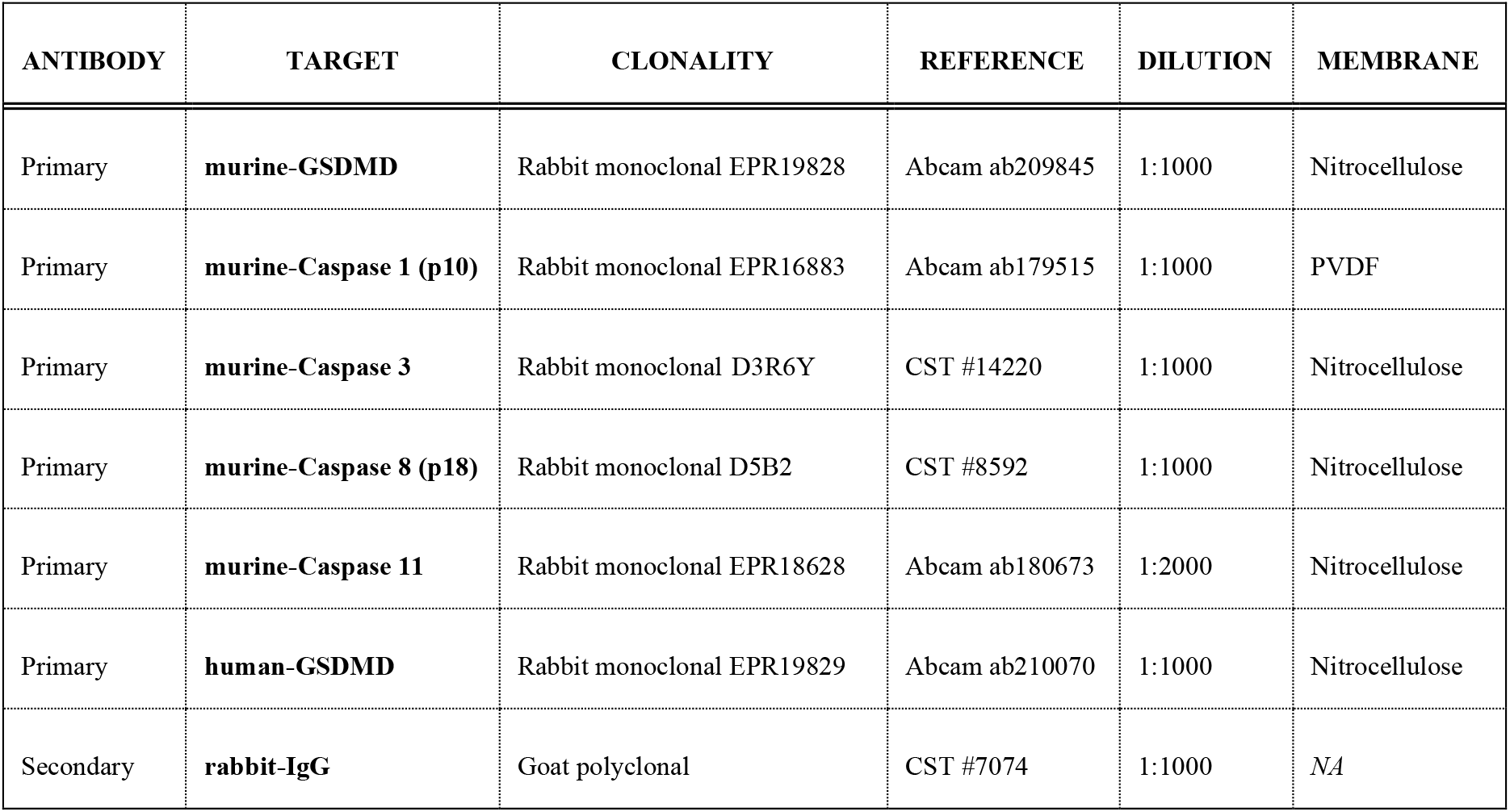
References of antibodies used for Western blot analyses.

| ANTIBODY | TARGET | CLONALITY | REFERENCE | DILUTION | MEMBRANE |
| --- | --- | --- | --- | --- | --- |
| Primary | **murine-GSDMD** | Rabbit monoclonal EPR19828 | Abcam ab209845 | 1:1000 | Nitrocellulose |
| Primary | **murine-Caspase 1 (p10)** | Rabbit monoclonal EPR16883 | Abcam ab179515 | 1:1000 | PVDF |
| Primary | **murine-Caspase 3** | Rabbit monoclonal D3R6Y | CST #14220 | 1:1000 | Nitrocellulose |
| Primary | **murine-Caspase 8 (p18)** | Rabbit monoclonal D5B2 | CST #8592 | 1:1000 | Nitrocellulose |
| Primary | **murine-Caspase 11** | Rabbit monoclonal EPR18628 | Abcam ab180673 | 1:2000 | Nitrocellulose |
| Primary | **human-GSDMD** | Rabbit monoclonal EPR19829 | Abcam ab210070 | 1:1000 | Nitrocellulose |
| Secondary | **rabbit-IgG** | Goat polyclonal | CST #7074 | 1:1000 | NA |

### Native gel electrophoresis and Western blots (WB)

RAW-ASC cells were collected 12 h post-transfection by removal of the medium and direct lysis using commercial lysis buffer maintaining caspase integrity and activity (from caspase 3/7 activity kit): 20 mM Tris-HCl pH 7.5, 150 mM NaCl, 1 mM Na_2_ EDTA, 1 mM EGTA, 1% v/v Triton X-100, 20 mM sodium pyrophosphate, 25 mM sodium fluoride, 1 mM β-glycerophosphate, 1 mM Na3VO4, 1 μg/ml leupeptin (CST), for 15 min on ice. After centrifugation for 10 min at full speed, proteins were recovered in the supernatant and protein concentration was determined and adjusted by Bradoford dosage (BioRad). Native protein sample buffer was then added, and electrophoresis was performed on polyacrylamide 4-15% gradient gel in 1X Tris-Glycine native buffer, at 90 V for 2 h. Transfers and Western blots were performed as previously described after denaturing SDS-PAGE.

### LPS-Caspase 11 binding assay

Interaction between recombinant caspase 11 (Enzo) and *E. coli* LPS (Invivogen) or *L. interrogans* LPS from serovar Manilae strain L495 was addressed by binding assays performed by incubation of 100 ng of recombinant enzyme with 50-200 ng of LPS in PBS for 1 h at 37°C. Sample were then loaded directly on home-casted native 4.5% (stacking) / 20% (running) highly reticulated polyacrylamide gels, that do not allow the entrance of LPS and bound material and only allow migration of unbound proteins. Electrophoresis was performed in 1X Tris-Glycine native buffer, at maximum 40V for minimum 6 on ice to avoid gel heating. Transfers and Western blots were performed as previously described after denaturing SDS-PAGE.

### *In vivo* infection with *Leptospira*

Adult C57BL/6J mice WT were obtained from Janvier Labs whereas IL1Receptor knock-out (ko) mice were bred at Institut Pasteur Paris animal facility. For each infection, groups of *n*=3 mice were infected *via* the intra-peritoneal (IP) route with either sublethal (2 x 10^6^ bacteria/mouse) or lethal (2 x 10^8^ bacteria/mouse) doses of *L. interrogans* serovar Copenhageni strain Fiocruz L1-130, in 200 μL of endotoxin-free PBS. Naive animals were injected with 200 μL of PBS alone. At day 3 or day 15 post-infection, animals were euthanized by cervical dislocation and organs were recovered and frozen at −80°C. Kidneys at day 3 were used for RT-qPCR analyses of cytokines mRNA and kidneys at day 15 were used for qPCR determination of bacterial loads.

### Ethic statement on animal use

All experiments performed on animals were conducted in accordance with the Animal Care guidelines and following the European Union Directive 2010/63 EU. Protocols were all approved beforehand (#2013-0034 and HA-0036) by the ethic committee of the Pasteur Institute, Paris, France (CETEA#89), in compliance with the French and European regulations on animal welfare and according to the Public Health Service recommendations.

### mRNA analyses by RT-qPCR

mRNA was extracted either from the frozen kidneys recovered 3 days post-infection, or from frozen mouse BMDMs using RNeasy Mini kit (Qiagen). Reverse transcription (RT) was performed using Superscript II reverse transcriptase (Invitrogen) according to the manufacturer’s recommendations. Generated cDNAs were used for quantitative PCR (qPCR) on a StepOne Plus real-time PCR machine (Applied Biosystems), with primers and probes targeting murine HPRT, caspase 11, IL1 β, IL6 and RANTES (**Table 2**). The following settings were used (relative quantification program): 50°C for 2 min, followed by 95°C for 10 min and by 40 cycles of 95°C for 15 s and 60°C for 1 min. Data were analyzed using the comparative 2^-ΔΔCt^ method, using a first normalization by internal control HPRT and a second normalization by non-infected controls.

**Table 2.**
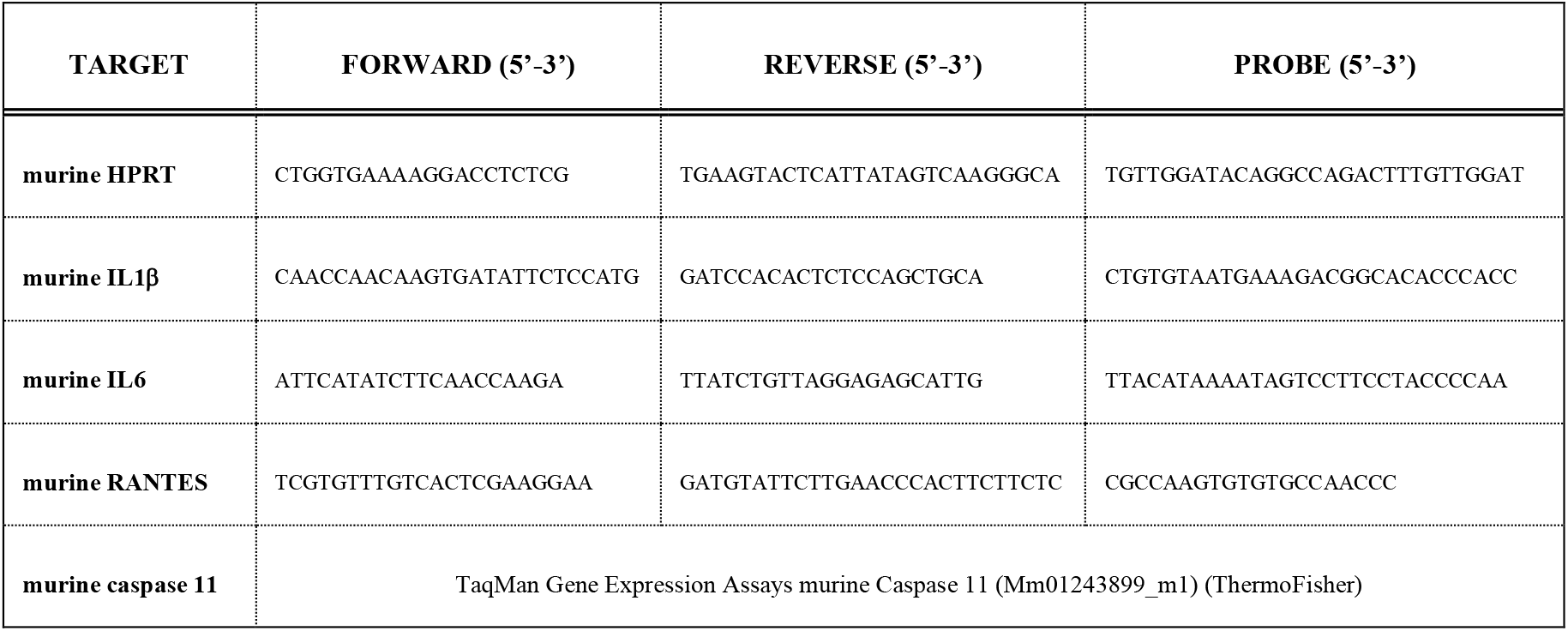
Primers and probes used for RT-qPCR analyses.

### Bacterial loads determination by qPCR in kidneys

Leptospiral DNA was extracted from the frozen kidneys recovered 15 days post-infection. Kidneys were mechanically disrupted with metal beads during 3 min at 4°C and DNA was then extracted using QIAmp DNA kit (Qiagen). Leptospiral DNA was specifically targeted using primers and probes designed in the *lpxA* gene (*L. interrogans* serovar Copenhageni strain Fiocruz L1-130): (Forward) 5,-TTTTGCGTTTATTTCGGGACTT-3’; (Reverse) 5’-CAACCATTGAGTAATCTCCGACAA-3’; (Probe) 5’-TGCTGTACATCAGTTTTG-3’ (Holzapfel *et al*., 2020). Normalization was performed using *nidogen* gene and quantitative PCR (qPCR) was performed on a StepOne Plus real-time PCR machine (Applied Biosystems), with the following settings (absolute quantification program): 50°C for 2 min, followed by 95°C for 10 min and by 40 cycles of 95°C for 15 s and 60°C for 1 min.

### Statistical analyses

All statistical analyses were performed using Student’s *t*-test with corresponding *p* values: * for *p* < 0.05; ** for *p* < 0.01 and *** for *p* < 0.001.

## Acknowledgments

We acknowledge Damien Arnoult, Stephen Girardin and Dana Philpott for fruitful discussions and for their kind suggestions. We also acknowledge Sonia Lacroix-Lamandé and Sylvain Bourgeais (UEPAO, INRAE CVL, France) as well as Nadia Benaroudj (Institut Pasteur Paris, France) for kindly providing the bovine and hamster bones, respectively. Finally, we acknowledge Frédérique Vernel-Pauillac for initial training of students.

